# Experimental Granulomatous Pulmonary Nocardiosis in Balb/c Mice

**DOI:** 10.1101/055293

**Authors:** Roque M. Mifuji Lira, Alberto Yairh Limón Flores, Mario César Salinas Carmona, Alejandro Ortiz Stern

## Abstract

Pulmonary nocardiosis is a granulomatous disease with high mortality that affects both immunosuppressed and immunocompetent patients. The mechanisms leading to the establishment and progression of the infection are currently unknown. An animal model to study these mechanisms is sorely needed. We report the first in vivo model of granulomatous pulmonary nocardiosis that closely resembles human pathology. BALB/c mice infected intranasally with two different doses of GFP-expressing *Nocardia brasiliensis* ATCC700358 (NbGFP), develop weight loss and pulmonary granulomas. Mice infected with 10^9^ CFUs progressed towards death within a week while mice infected with 10^8^ CFUs died after five to six months. Histological examination of the lungs revealed that both the higher and lower doses of NbGFP induced granulomas with NbGFP clearly identifiable at the center of the lesions. Mice exposed to 10^8^ CFUs and subsequently to 10^9^ CFUs were not protected against disease severity but had less granulomas suggesting some degree of protection. Attempts to identify a cellular target for the infection were unsuccessful but we found that bacterial microcolonies in the suspension used to infect mice were responsible for the establishment of the disease. Small microcolonies of NbGFP, incompatible with nocardial doubling times starting from unicellular organisms, were identified in the lung as early as six hours after infection. Mice infected with highly purified unicellular preparations of NbGFP did not develop granulomas despite showing weight loss. Finally, intranasal delivery of nocardial microcolonies was enough for mice to develop granulomas with minimal weight loss. Taken together these results show that Nocardia brasiliensis microcolonies are both necessary and sufficient for the development of granulomatous pulmonary nocardiosis in mice.

## Introduction

Pulmonary nocardiosis is a serious disease with high mortality [1]. While it usually affects immunocompromised patients there are reports of the disease in apparently normal hosts [2–4].

Classical bacteriological studies had shown that most cases of pulmonary nocardiosis were due to *Nocardia asteroides* but advances in genomic taxonomy have shown these to be mostly *N. farcinica, N. nova* and *N. cyriacigeorgica* [5], while most nocardial infections of the skin are caused by *Nocardia brasiliensis* [6]. Work by Beaman et. al. spanning almost three decades studied nocardial infections in a wide variety of animal hosts using *Nocardia asteroides* and other species of nocardia. Studies with *Nocardia asteroides* showed that mice were able to successfully control pulmonary infection within a week [7], through the coordinated action of IFNy producing neutrophils [8], recruited via CXCR2 [9], TyS cells and IL–17 cells [10]. Additionally, work by Abraham et. al. [11] inoculating *N. brasiliensis* intravenously to mice showed abscess formation in the lungs. No other successful models of pulmonary nocardiosis have been reported, even when rabbit lung macrophages have been succesfully infected by nocardia *in vitro* [12]. Pulmonary nocardiosis in humans however, usually presents as a chronic or subacute supurative disease with formation of granulomas and consolidated areas of the lung [2]. Hence, a proper pulmonary granulomatous nocardiosis model is sorely lacking.

Unlike pulmonary nocardiosis, an adequate murine model of actynomicetoma exists. Mice infected with 10^7^ CFUs of *Nocardia brasiliensis* in the footpad develop a progressive disease with formation of abscesses and fistulae characteristic of human pathology [13]. Working up from the mycetoma model and through the use of GFP-expressing *Nocardia brasiliensis* (NbGFP) (to facilitate detection) [14] we sought to develop a new model of pulmonary nocardiosis to better study the disease and its immunological aspects. We report the successful development of an experimental pulmonary granulomatous nocardiosis model that should be useful in many settings: to study lung immune responses towards nocardia, to investigate granuloma formation through the use of an infectious agent of minimal virulence to humans and in pre-clinical trials of new anti-nocardial agents.

## Results

### Intranasal instillation of *Nocardia brasiliensis* leads to pulmonary granulomas

NbGFP isolated from a murine mycetoma and tested for GFP positivity using a photodocumenter set to λ 488 nm was grown for 72 hrs on BHI medium without stirring. After 72 hrs NbGFP formed a biofilm on the surface of the culture media (Sup Fig. 1). Biomass was washed with saline solution and homogenized with a Potter-Elvehjem homogenizer. The bacterial suspension was left to settle and the top half, containing a homogeneous unicellular suspension (as evaluated with a Neubauer chamber) was adjusted to a concentration of 10^10^ CFUs per 50 μl. Serial 1/10 dilutions were then prepared to infect mice intranasally. These same serial 1/10 dilutions were plated and CFUs were corroborated by colony counting.

Based on previous infectious models using *Nocardia spp.* [10] we first tested a dose of 10^6^ CFUs without any infected BALB/c mice showing symptoms or signs of disease. In a second experiment mice receiving 10^10^ CFUs died within 24 hours and were not studied any further. Mice were then tested in two independent experiments with doses of 10^9^ and 10^8^ CFUs. Most mice receiving 10^9^ CFUs died or had to be sacrificed within a week after losing in average up to 21.5% of their original body weight by day 2 (p= 6.45E–008 vs day 0) without showing signs of recovery (Fig 1A and B). On the contrary 80% of mice receiving 10^8^ CFUs survived the infection after mild weight loss (8.4% day 2 vs day 0 p= 0.0001) that was completely recovered within three weeks (Fig 1A and B). After sacrifice, lungs were excised and filled with a buffered 10% formalin solution. Images of the lungs were captured with a photodocumenter at λ 488 nm and then further processed for histological analysis. Lungs of mice infected with 10^9^ CFUs had high numbers of GFP positive dots (Fig 1C) shown to be Kinyoun-positive granules upon histological examination (Fig 1E). Lungs of mice infected with 10^8^ CFUs analysed 21 days after infection also had GFP positive dots, albeit at much lower numbers (white arrows in Fig 1D). These lesions appear as granulomas containing Kinyoun-positive nocardia (Fig 1F left panel). Mature granulomas were defined by the clear presence of activated macrophages and lymphocytes surrounding the nocardia (Fig 1F right panel). We decided to follow the long term progression of the disease in two mice infected with low doses of NbGFP. Six months after the infection both mice looked unhealthy and had difficulty moving. Mice were euthanized and lungs processed as previously described. Lungs of these mice had massive abscesses and upon histological examination these were shown to contain a large amount of cellular debris, nocardial microcolonies and severe disruption of surrounding tissue (Fig 1G).

**Figure 1:**
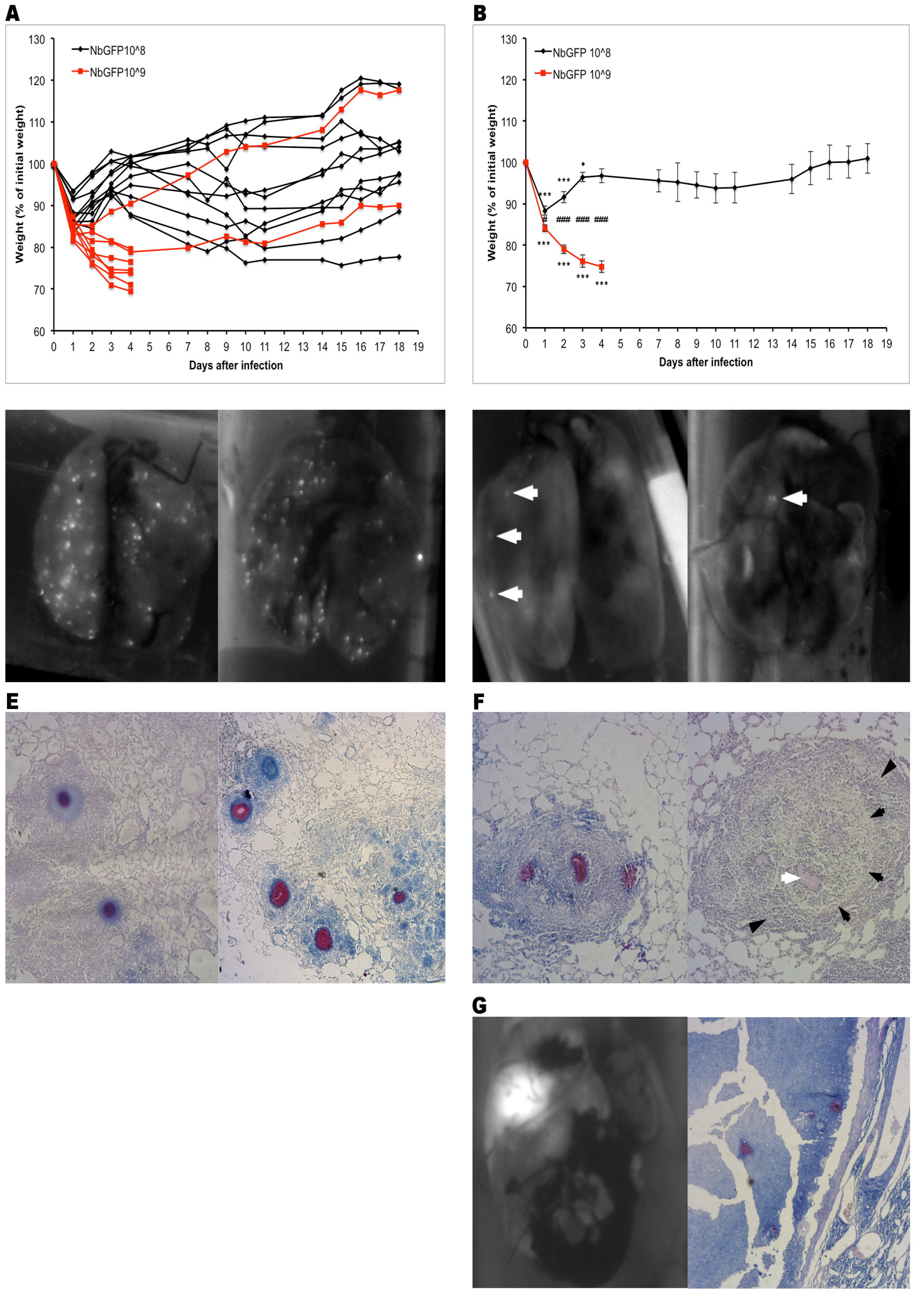
**Intranasal instillation of *Nocardia brasiliensis* leads to pulmonary granulomas.** A) BALB/c mice received 10^8^ (black) or 10^9^ (red) NbGFP CFUs and were weighted daily. Individual mice are shown B) Data from A analysed as average +/− s.e.m. The two surviving 10^9^ CFUs-receiving mice were excluded. C) Lungs from mice receiving 10^9^ CFUs were analysed at day 4 post infection by whole lung imaging at λ 488 nm. Representative images of two lungs are shown. D) Lungs from mice receiving 10^8^ CFUs were analysed at day 21 post infection by whole lung imaging at λ 488 nm. Representative images of two lungs are shown. White arrows point to GFP+ lesions. E) Representative images of lesions present in mice receiving 10^9^ CFUs at day 4 post infection. Kinyoun's stain. Red stained NbGFP is evident at the core of the lesion. F)Representative images of lesions present in mice receiving 10^8^ CFUs at day 21 post infection. Left panel: Kinyoun's stain. Red stained NbGFP is evident at the core of the lesion. Right panel: H&E staining. White arrow: NbGFP, black arrows: activated macrophages, black arrowheads: lymphocytes. G) Whole lung imaging and histological (Kinyoun's stain) analysis of a lung from a mice six months after infection with 10^8^ NbGFP CFUs. Graphs show data from two independent experiments (n=11 (10^8^), 8 (10^9^)) * p<0.05 vs. initial weight, ** p<0.01 vs. initial weight, *** p<0.001 vs. initial weight, # p<0.05 10^8^ vs. 10^9^, ## p<0.01 10^8^ vs. 10^9^, ### p<0.001 10^8^ vs. 10^9^.

### 10^8^ dose intranasal infection diminishes granuloma load upon 10^9^ dose challenge

Even though mice that received 10^8^ CFUs did not completely clear the infection, as evidenced by the presence of granulomas by three weeks and abscesses six months after inoculation, we sought to test if exposure to the lower doses of NbGFP provided protection against a higher dose challenge. Two groups of mice received either saline solution or 10^8^ CFUs NbGFP and 21 days afterwards both groups received 10^9^ CFUs. Both groups lost weight similarly after the second challenge (17.2 +/− 4% vs 13.3 +/− 0.09% p=0.52 low dose vs saline, respectively) and surprisingly mice in both groups survived with the exception of a single mouse in the saline receiving group (Fig 2A). However upon examination of the lungs in the photodocumenter, lungs from mice that had received a previous 10^8^ dose of NbGFP had significantly less dots than mice that had received saline solution (Fig 2B and C). We hypothesized that we could have inadvertently given mice a slightly smaller dose than 10^9^ during the second challenge or alternatively that mice in our facility could have had a concomitant non-diagnosed infection that altered the course of the disease. We then repeated the experiment with a new set of SPF mice. Just like in the first experiment, both groups of mice, those that received 10^8^ CFUs and those that received saline solution lost weight similarly upon challenge with 10^9^ CFUs. However disease severity was greater than in previous experiments and all mice had to be sacrificed 4 days after the infection (Fig 2D). Upon analysis of the lungs with the photodocumenter, mice that had been exposed to a lower dose prior to challenge had reduced numbers of granulomas when compared with naive mice exposed to saline solution (Fig 2E and F). Taken together these experiments suggest that severity of the disease during the first days after infection may not be directly related with granuloma burden and that previous exposure to 10^8^ CFUs of NbGFP protects mice against developing higher number of lesions upon exposure to 10^9^ CFUs.

**Figure 2:**
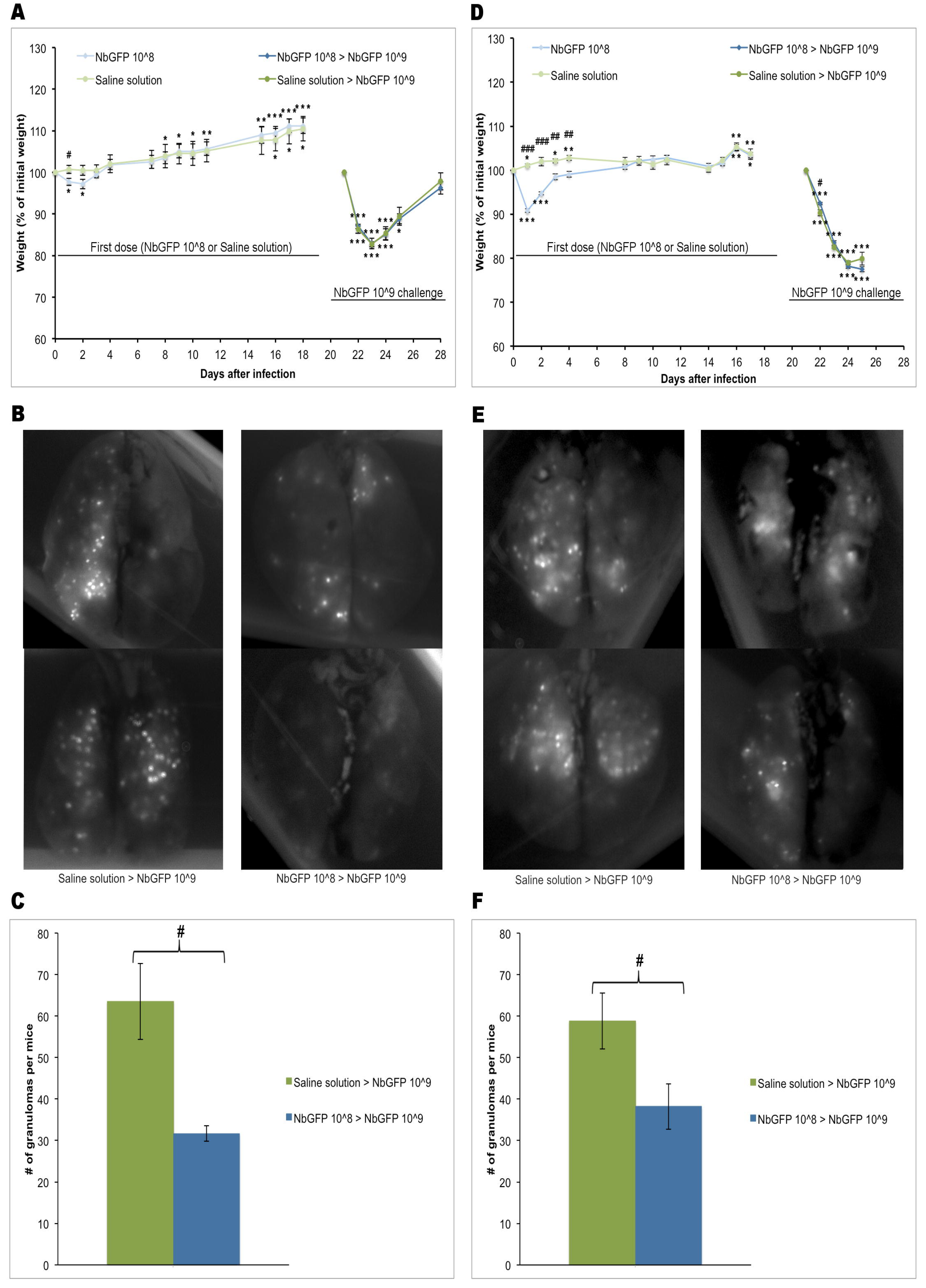
**Low dose intranasal infection diminishes granuloma load upon high dose challenge.** A) BALB/c mice received saline (green) or 10^8^ (blue) NbGFP CFUs and were weighted daily. At day 21 post infection mice were challenged with 10^9^ NbGFP CFUs. Weight at day 21 was re-normalized as 100%. B) Lungs from mice challenged with 10^9^ CFUs at day 21 were analysed at day 7 post challenge by whole lung imaging at λ 488 nm. Representative images of two lungs per group are shown. C) Average number of lesions per animal from experiment depicted in A and B (n=5 mice per group) D) SPF BALB/c mice received saline (green) or 10^8^ (blue) NbGFP CFUs and were weighted daily. At day 21 post infection mice were challenged with 10^9^ NbGFP CFUs. Weight at day 21 was re-normalized as 100% E) Lungs from SPF/mice challenged with 10^9^ CFUs at day 21 were analysed at day 4 post challenge by whole lung imaging at λ 488 nm. Representative images of two lungs per group are shown. F) Average number of lesions per animal from experiment depicted in D and E (n= mice per group). Graphs show average +/− s.e.m. (n=5 per group) * p<0.05 vs. initial weight, ** p<0.01 vs. initial weight, *** p<0.001 vs. initial weight, # p<0.05 10^8^ vs. 10^9^, ## p<0.01 10^8^ vs. 10^9^, ### p<0.001 10^8^ vs. 10^9^.

### Heat killed NbGFP causes weight loss upon intranasal delivery

Since granuloma burden and disease severity did not seem to have an absolute correlation, we wanted to know if bacterial components were capable of causing weight loss in BALB/c mice regardless of granuloma formation. Mice exposed to 10^9^ heat-killed CFUs lost up to 9.4 % of their body weight one day after inoculation (Sup Fig 2). Image analysis of the lungs of those mice revealed that they did not develop granulomas at any time point (Sup Fig 2).

### No cell could be identified as a target for NbGFP in the lung

Current models of nocardiosis pose that the macrophage is the prime target for nocardial infections [12,15,16]. In order to test if alveolar macrophages were the target for NbGFP, we isolated macrophages from bronchoalveolar lavages (BAL) of naive mice and exposed them to NbGFP (1:10, Macrophage:NbGFP) for up to 48 hours and imaged the cells every 24 hours. Unlike bone marrow derived macrophages (Fig 3A), alveolar macrophages were not susceptible to infection by NbGFP (Fig 3A). Additionally alveolar macrophages isolated from BAL of inoculated mice were also not infected (Fig 3A). We then hypothesized that it was possible for other phagocytic cells in the airway to be subverted by NbGFP. Given the fact that after inoculation of 10^9^ CFUs only some 60 lesions are apparent in the lungs of infected mice, such cell should be found in very low numbers. We isolated BAL from infected mice 24 and 48 hours after infection to analyse it by flow cytometry targeting eosinophils as a low prevalence possible target for NbGFP (Fig 3B). BAL analysis showed that eosinophils while present in the lungs of infected animals, did not appear infected (Fig 3B). Flow cytometry showed that most GFP positive cells appeared to be neutrophils, however, by 48 hours the number of GFP+ neutrophils had sharply declined, suggesting addecuate killing of phagocytosed nocardia as reported by Beaman [9]. Additionally, the number of GFP+ neutrophils was not compatible with the number of lesions present in the lung 7 days after the infection suggesting that neutrophils are not the target of the infection (Fig 3C).

**Figure 3:**
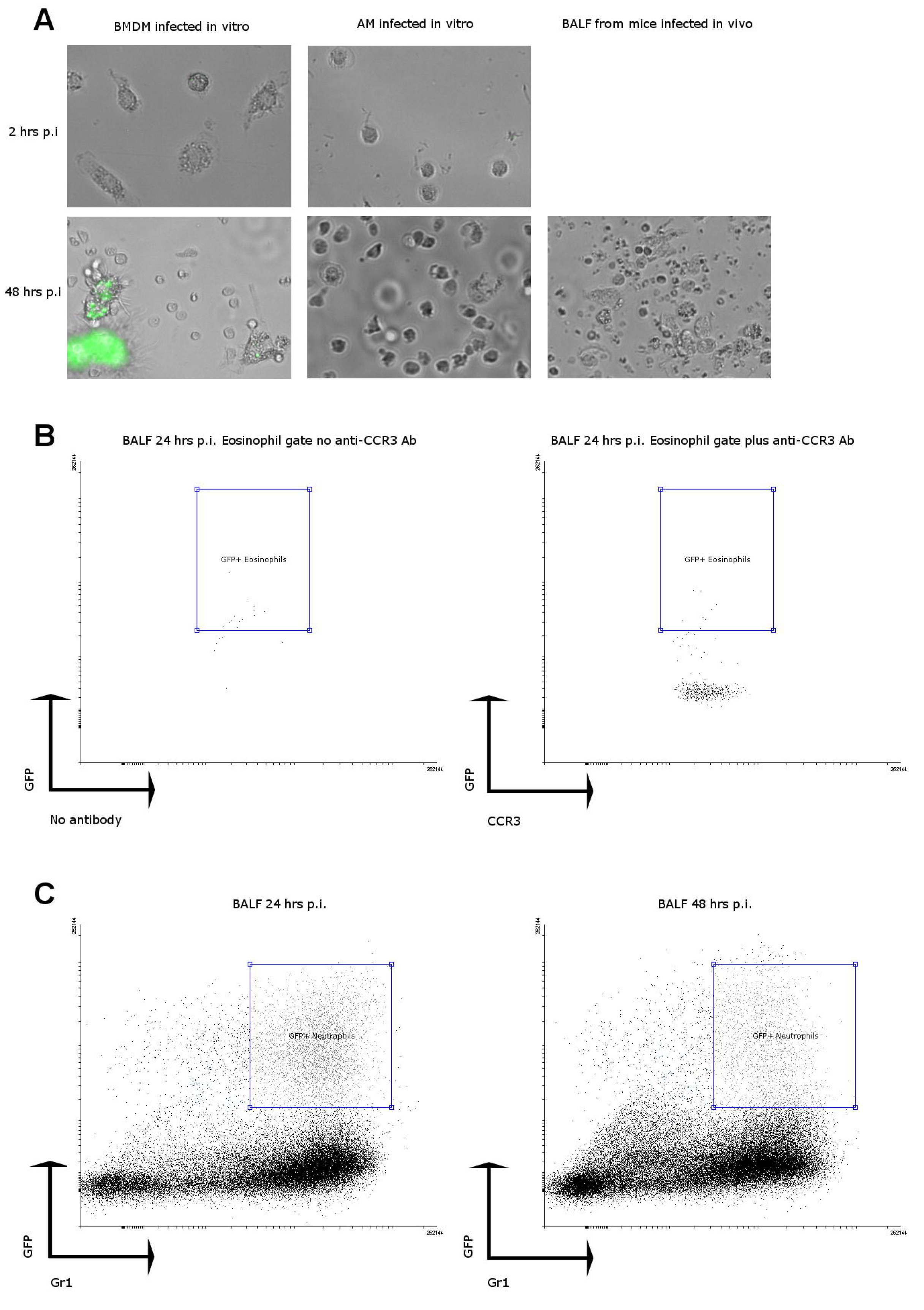
Neither alveolar macrophages nor eosinophils nor neutrophils are targets for NbGFP infection in the lung. A) Bone marrow derived macrophages (BMDM) or alveolar macrophages (AM) were incubated *in vitro* with NbGFP (1:10) and imaged at 2 hours and at 48 hrs post infection. Cells from BALF of infected mice were imaged 48 hrs after infection to look for infected alveolar macrophages. Representative images are shown. B) BALF analysis of eosinophils by flow cytometry. Left panel shows non-specific events detected within the eosinophil gate in the absence of CCR3 staining. Right panel shows no increase in GFP+ events amongst eosinophils stained with an anti-CCR3 antibody. C) BALF analysis of granulocytes by flow cytometry at 24 (left panel) and 48 (right panel) hours post infection.

### Microscopic bacterial microcolonies are responsible for the formation of granulomas

We then turned to confocal microscopy to try to identify cells that could harness large numbers of bacteria at early time points. Analysis of lungs at 6 hrs, and 24 hrs after infection did not reveal the presence of spots detectable with the photodocumenter while granulomas were clearly present at 7 days (Fig 4A). Microscopic confocal analysis of lung sections from mice infected with 10^9^ CFUs NbGFP at 6 and 24 hours after infection revealed the presence of microscopic microcolonies of NbGFP surrounded by inflammatory cells (Fig 4B), but no microcolonies were found inside cells of any kind. Growth curves for NbGFP suggest a doubling time of 7.8 hours (Fig 4C), which is incompatible with the presence of bacterial microcolonies at such early time points. Infection with wild type *Nocardia brasiliensis* (ATCC700358) also produced granulomas when instilled intranasally (Sup Fig.3).

**Figure 4:**
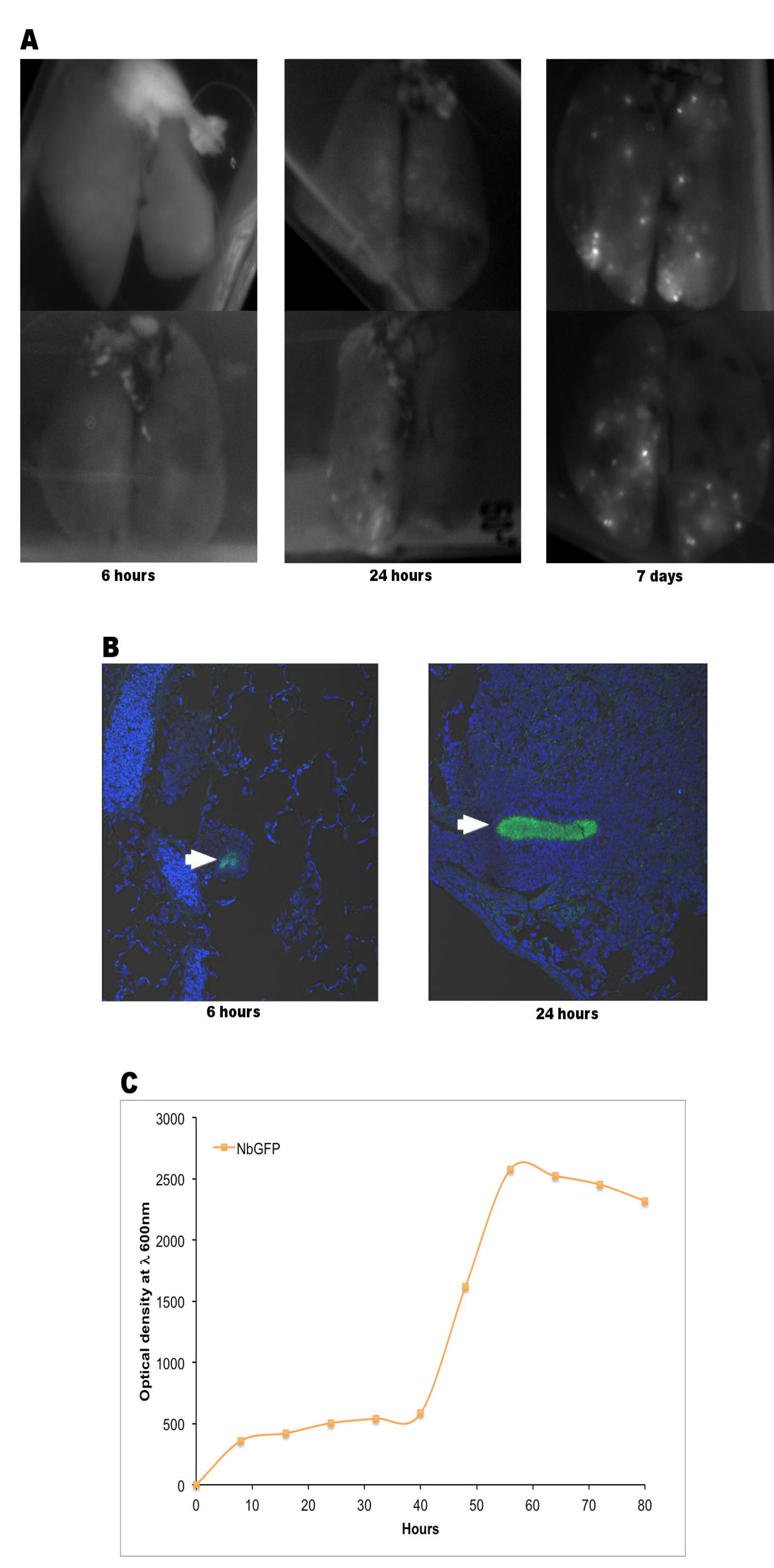
NbGFP microcolonies are present in the lungs of infected mice as early as 6 hours post infection. A) Lungs from mice receiving 10^9^ CFUs were analysed at 6 hours, 1 day and 4 days post infection by whole lung imaging at λ 488 nm. Representative images of two lungs per time point are shown. B) Confocal microscopy images of lung sections at 6 and 24 hours post infection. C) Growth curve of NbGFP under constant stirring to prevent biofilm formation.

To test if bacterial microcolonies were responsible for granuloma formation we removed NbGFP microcolonies from the suspension prior to intranasal instillation. Mice that received a highly purified unicellular 10^9^ CFUs dose, lost almost as much weight as mice receiving a regular 10^9^ CFUs suspension, however none of the mice developed lesions 4 days after inoculation while mice that received the regular suspension developed granulomas (Fig 5A, B and E). Finally we isolated bacterial microcolonies and prepared a suspension that was examined visually to contain between 50 to 100 microcolonies per 50 μl. Mice infected with this suspension lost very little weight which they recovered at day 2. Upon both image and microscopic analysis these mice were shown to have nocardial lesions by day 4 (Fig 5D and F). Mice infected with bacterial microcolonies and analysed 21 days after infection also had lesions corresponding to mature granulomas as evidenced by the presence of activated macrophages and lymphocytes surrounding the nocardia (Fig 5G).

**Figure 5:**
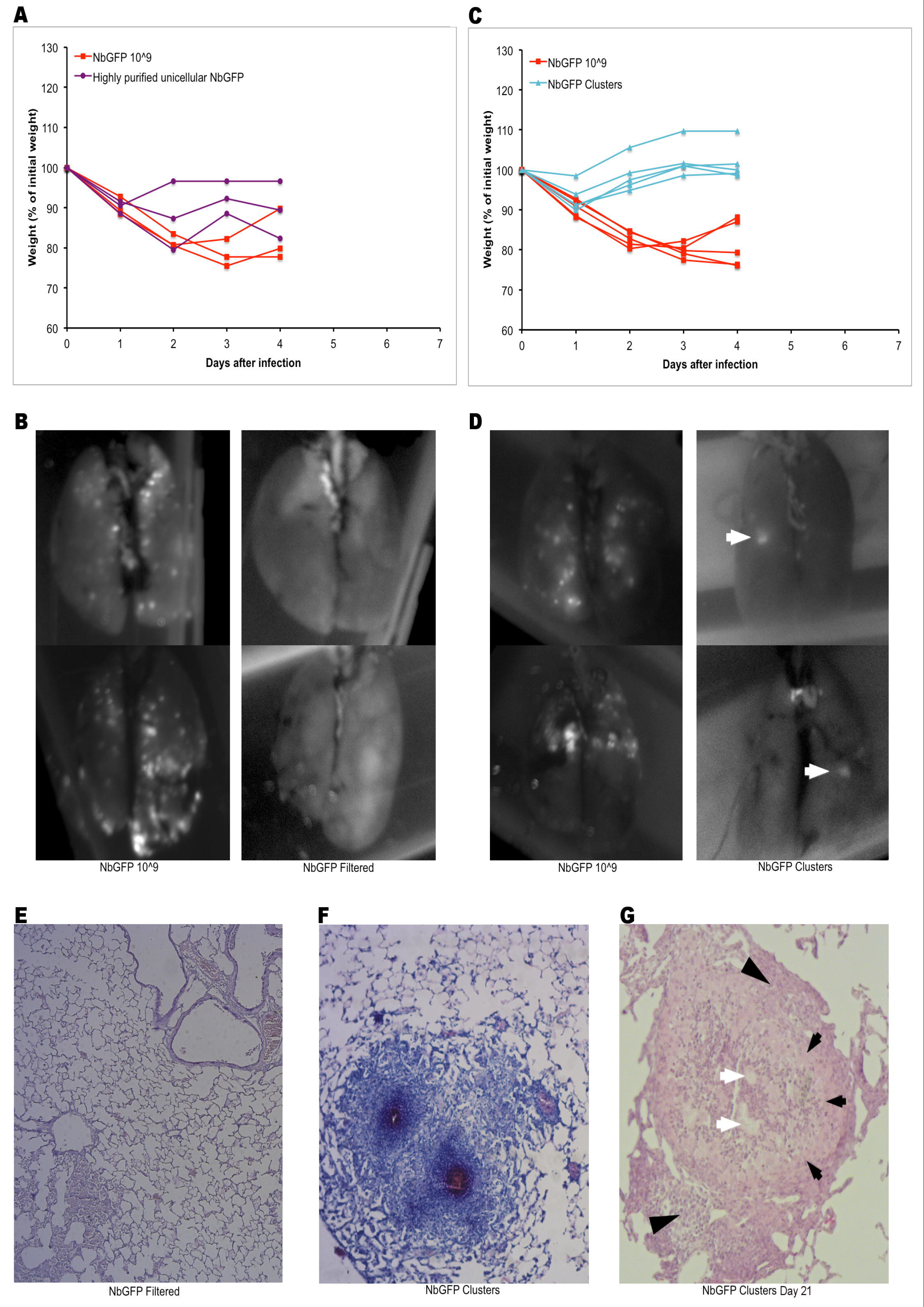
Microscopic bacterial microcolonies are responsible for the formation of granulomas. A) BALB/c mice (n=3 per group) received 10^9^ (red) or 10^9 Hlghly purified single-cen^ (purple) NbGFP CFUs and were weighted daily. B) Lungs from mice in A were analysed at day 4 post infection by whole lung imaging at λ 488 nm. Representative images of two lungs per group are shown. C) BALB/c mice (n=5 per group) received 10^9^ (red) CFUs or ≈100 (teal) NbGFP microcolonies and were weighted daily. D) Lungs from mice in D were analysed at day 4 post infection by whole lung imaging at λ 488 nm. Representative images of two lungs per group are shown. White arrows point to GFP^+^ lesions. E) Representative histology of lungs in mice receiving 10^9 Highly purified single-ceU^ CFUs at day 4 post infection. Kinyoun's stain. F) Representative images of lesions present in mice receiving microcolonies. Kinyoun's stain. Red stained NbGFP is evident at the core of the lesion. G) Representative H&E image of lesions present in mice 21 days after infection with purified microcolonies. White arrows: NbGFP, black arrows: activated macrophages, black arrowheads: lymphocytes.

### Inflammation drives lethality of infection with 10^9^ CFUs

While nocardial microcolonies drive the formation of granulomas, the mechanism for the lethality seen when infecting mice with 10^9^ CFUs was not clear. The acute weight loss observed in these mice point towards an exacerbated inflammatory response. To evaluate the inflammatory response we analysed BAL cell numbers, protein content and IL–1β levels of mice infected with either 10^8^ or 109 CFUs 24 and 48 hours after infection (Sup Fig. 4). BAL cell numbers at 24 hours post infection were not different between the 10^9^ or 10^8^ dose. However, the amount of GFP+ neutrophils was, by far, larger on animals infected with the higher dose. At 48 hours post-infection BAL cell numbers and neutrophil content was clearly diminishing on mice receiving 10^8^ CFUs. In clear contrast, BAL cell numbers and total neutrophils increased between 24 and 48 hours in mice given 10^9^ CFUs. Additionally, the number of GFP+ neutrophils remained elevated (Sup Fig. 4B–F). BAL protein levels were significantly elevated only in mice receiving 10^9^ CFUs both at 24 and 48 hours post infection (Sup Fig. 4G). IL–1μ levels were eight fold higher in mice receiving 10^9^ CFUs (148.9 ± 21.5 pg/ml) compared with mice receiving 10^8^ CFUs (18.3 ± 5.6 pg/ml) 24 hours post infection. Furthermore, IL–1β remained elevated at 48 hours post infection in mice receiving 10^9^ CFUs (100.3 ± 13.02 pg/ml) while they became undetectable in mice receiving 10^8^ CFUs (Sup Fig. 4H).

Taken together these results provide very strong evidence that microscopic bacterial microcolonies are both necessary and sufficient for *Nocardia brasiliensis* to cause granulomatous pulmonary nocardiosis in mice.

## Discussion

To our knowledge this is the first description of an animal model of pulmonary granulomatous nocardiosis. While work by Beaman and collaborators has helped to understand some of the immune responses towards nocardial infections, much has been missed by the fact that animals infected intranasally with *Nocardia asteroides* do not develop granulomatous disease [10]. Previous work by Abraham and collaborators showed that mice receiving intravenous *N. brasiliensis* developed lung abscesses containing the pathogen but not granulomas. This discrepancy with our results could be explained by the various methodological differences, namely, the route of administration, bacterial load, bacterial strain and the use of steroids [11].

In pursuing the goal of developing a granulomatous model we departed from the methodology used by Beaman [17]. The main difference, apart from our use of *Nocardia brasiliensis,* is that our inoculum is derived from a disrupted biofilm instead of a culture grown in agitation. While this methodology, the disruption of *N. brasiliensis* biofilms, had been reported to produce single organisms [13] we found that disruption is not complete and microcolonies remain in bacterial suspensions prepared this way. These microcolonies were not easily identified when our first set of experiments were performed, the most likely explanation as to why we did not detect them could be the high dilution needed to adjust CFUs for infection which precluded their detection (Sup Fig. 1). Thus the inoculum used to infect mice in our first experiments was a mixture of single-celled organisms and bacterial microcolonies. Mice infected with this preparation still presented a dose/response relationship; mice infected with higher CFUs evolved with worst clinical outcomes and more granulomas. Both our experiments to test for protection and those aimed at identifying the target cell of the infection were carried out under the idea that single cell organisms were responsible for the infection.

The finding that alveolar macrophages were not susceptible to *Nocardia brasiliensis* infection neither *in vitro* nor *in vivo* was very unexpected in the light of current knowledge of nocardial diseases [15] and prompted us to look for alternatives. Analysis of lungs at very early time points after infection were crucial to understanding the pathogenesis of the disease since presence of microcolonies as early as six hours post infection were completely incompatible with single-cell organisms being responsible due to nocardial doubling times of 8 hours. It will be important to understand the role of alveolar macrophages within the context of nocardial granulomas to see if their organization is comparable to that of other granulomatous diseases (i.e. tuberculosis) where pro-inflammatory macrophages lie mostly at the center and anti-inflammatory macrophages reside in the periphery of the granuloma [18]. Analysis of mice that received 10^8^ CFUs of the unfiltered NbGFP or purified microcolonies 21 days after the infection revealed that the lesions seen at 4 days indeed progress towards the formation of a mature granuloma with activated macrophages and lymphocytes surrounding the nocardia. Studying the phenotype of these cells should yield rich results regarding granuloma formation and progression.

It is interesting to notice that granulomas generated in the lung in response to *N.* brasiliensis nocardial microcolonies very much resemble the lesions seen in subcutaneous actinomycetomas.While working on this manuscript we conducted an extensive image search looking for histolopathological samples of human pulmonary nocardiosis due to *Nocardia brasiliensis* to directly compare with our findings but we were unsuccessful, none seem to be documented. However, one report of an invasive *N. brasiliensis* infection of the skin with pulmonary involvement showed lesions extremely similar to the ones we show in this manuscript [19]. Additionally, granulomas due to other *Nocardia spp.* present themselves with a similar architecture to the lesions we describe in this manuscript, namely, extracellular bacteria surrounded by macrophages and lymphocytes [20]. The importance of granulomas in pulmonary nocardiosis is further underscored by the fact that pulmonary nodules are the second most common radiographic finding in pulmonary nocardiosis [21]. Furthermore in a report of endobronchial nocardiosis masses of nocardia, very much like our microcolonies were seen upon biopsy of the lesions [22].

Our experiments with highly purified single-cell organisms and microcolonies further clarified their role in the disease. Single-cell organisms, accounting for most of the 10^8^–10^9^ CFUs given to mice, appear responsible for the acute inflammatory response and weight loss seen upon infection. On the contrary, microcolonies did not cause such weight loss. While it could be possible to speculate that single-cell organisms could cause granulomas after longer time periods, we think this is unlikely, given that upon microscopic examination, no granulomatous or pre-granulomatous lesion could be seen four days after infection, a time at which colonies are already apparent to the naked eye upon culture in agar plates.

Our studies on protection suggested that weight loss was mostly due to an inflammatory response. Mice infected with 10^8^ CFUs and then re-challenged with 10^9^ CFUs lost as much weight as mice receiving saline and then a challenge with 10^9^ CFUs. When addressed directly it became clear that the inflammatory response to 10^9^ CFUs was of a grave magnitude comprising intense neutrophil recruitment, augmented protein leakage and high levels of IL1–β. Despite this inflammatory response, a role for immunological memory seems to be present in pre-challenged mice as evidenced by the reduced granuloma burden upon re-challenged. We are currently working in our lab to clarify the nature of this protection.

A current view of bacterial pathogenicity suggests that subversion of phagocytes is highly related with low infectious doses [23]. However this view assumes that single organisms, or multiple individual organisms, are responsible for infections. Our data adds one more possibility, the, ability to grow as microcolonies or to dislodge as microcolonies from biofilms. While biofilm formation has been largely explored as a problem on medical devices and other surfaces [24] the study of its role in other infectious routes has just started. microcolonies as infectious units of the airways should be limited by size, large microcolonies (>15 μm) would be unable to travel past the upper airways and microcolonies smaller than macrophages (< 8μm) should be effectively managed by professional phagocytes [25]. We are currently working towards pinpointing precisely the minimal microcolony size required for progression to granuloma as this will allow us to study granuloma formation and progression in both early and late phases of the disease.

In conclusion, *Nocardia brasiliensis* microcolonies are sufficient and necessary to cause granulomatous pulmonary nocardiosis in immunocompetent BALB/c mice. Additionally, this experimental model should be useful in helping to understand granulomatous diseases, through the use of an infectious agent of minimal virulence to humans, and in pre-clinical trials of new anti-nocardial agents.

## Materials and Methods

### Ethics Statement

All experimental protocols and care of the animals were approved by the Comití Institucional para el Cuidado y Uso de Animales de Laboratorio (CICUAL) within the Ethics Committee of the Facultad de Medicina, Universidad Autónoma de Nuevo León. The protocol registration with the 14 CICUAL Ethics Committee is: IN 13–005. The Ethics Committee, recognized by the National Bioethics Comission (C0NBI0ETICA19CEI00320130404) and the Secretaría de Salud via COFEPRIS (13CEI1903937), operates under national guidelines for the use, care and handling of laboratory animals as established by the Norma Oficial Mexicana (N0M–062–Z00–1999). Anesthesia was provided for all procedures involving animals using a Ketamine/Xylazine mixture.

### Inoculum preparation and growth analysis

Nocardia brasiliensis (ATCC 700 358) expressing green fluorescent protein (NbGFP) [14], was cultured in 50 ml BHI broth (Brain Heart Infusion, 0xoid, Hampshire, England) supplemented with gentamycin sulfate (Garamicina®, Schering-Plough S.A., Mexico) 26 μg/ml for 72 hours at 37° C without stirring. Biofilms from three Erlenmeyer flasks were resuspended in sterile saline and homogenized using a Potter-Elvehjem homogenizer to obtain a unicellular suspension. Suspensions were allowed to settle for 10 minutes and the top phase was recovered, centrifuged, weighted and resuspended to 250 mg/ml (which yields 10^10^ CFUs per 50 μl as corroborated by serial dilution plating in at least five independent experiments). From there different doses of colony-forming units (CFUs) were prepared through dilutions, corroborated using a Neubauer chamber and by serial dilution plating. For growth analysis NbGFP was grown under constant stirring and optical density sampled at A 600nm (SmartSpec Plus, Bio-Rad Laboratories) at shown intervals. Nocardial doubling times were calculated to be 7 hours and 28 minutes using exponential regression. Highly purified single-cell NbGFP inoculum was prepared by passing the nocardial suspension through a 4 μm pore filter. microcolonies were recovered from the filters and diluted to the desired concentration.

### Animals

Mice were housed at the animal facility of the Department of Immunology of the Facultad de Medicina, UANL under controlled temperature, humidity and lighting and *ad libitum* feeding. Male 6 to 8 weeks old, BALB/c and SPF BALB/c mice were anesthetized with an intraperitoneal injection of 200 μl of ketamine (Anesket®, PISA Agropecuaria, Mexico)/xylazine (Procin®, PISA Agropecuaria, Mexico) mixture, containing 2 mg and 0.4 mg respectively. Mice were infected with different doses of NbGFP via intranasal instillation (from 10^6^ to 10^10^ CFUs/mouse.) For the immune protection experiments, mice were instilled intranasally with sterile saline or NbGFP 10^8^ CFUs and 21 days later they were re-challenged with 10^9^ CFUs.

### Necropsy

Mice were sacrificed via cervical dislocation. Lsungs were excised, filled with, and submerged in a buffered 10% formalin solution.

### Macroscopic analysis

Inflated lungs were analysed with a photodocumenter (ChemiDoc™ MP System, Bio-Rad Laboratories), at λ 488nm. Images were analysed and adjusted for brightness and contrast using ImageJ. Lesions were manually scored by 2 independent researchers.

### Microscopic analysis

Inflated and fixed lungs in buffered 10% formalin solution for 48 hrs were cut and dehydrated in 70% ethanol for 2 hours, 80% ethanol 1 hour, 95% ethanol 1 hour, and 3 changes of 100% ethanol for 1 hour. After tissue dehydration, lungs were transferred to xylene (Xileno RA, CTR Scientific, Mexico) 3 changes of 1.5 hours each. Lungs were then submerged in 60° C paraffin (Paraplast ^®^ regular, Sigma-Aldrich, MO, U.S.A.) twice, for 2 hours, and embedded in individual cassettes. 5 μm lung sections were deparaffinized using 1.7% dishwashing soap solution as described by Amita Negi [26], and dyed using Kinyoun's stain (Carbol fuchsin, Sigma-Aldrich, MO, U.S.A., and Methylene Blue, Sigma-Aldrich, MO, U.S.A). Images were acquired using Carl Zeiss Axio Scope A1 equipped with AxioCam ERc 5s and imaging software AxioVision v. 4.8. For confocal microscopy, lung sections were deparaffinized and mounted in media with 4′,6–diamidino–2–phenylindole (DAPI) (VECTASHIELD^®^ with DAPI, Vector laboratories, CA, U.S.A). Images were acquired using Carl Zeiss LSM 710–NL0 Multiphoton Confocal Microscope equipped with imaging software ZEN (Zeiss efficient navigation) version 5.5.0.0.

### Bronchoalveolar lavage

Bronchoalveolar lavage fluid was obtained from naive or NbGFP infected mice (one or two days after infection) by 3x 1 ml washes with warm saline. Alveolar macrophages from naive mice were allowed to adhere to plastic culture plates for 6 hours before infection with NbGFP as described below for BMDM. Alveolar macrophages from infected mice were incubated for 48 hrs and imaged as described for BMDM.

### Flow Cytometry

BAL cells from infected mice were incubated with 1/500 dilutions of A647–antiCCR3 (clone J073E5, Biolegend), PE-Gr1 (clone RB6–8C5, Biolegend) for 30 minutes at room temperature and washed 3x in 10 volumes of 0.5% BSA PBS. Cells were processed using a LSRFortessa (BDBiosciences) equipped with Diva software. Analysis and figure preparation was performed with Flowing Sofware 2 (http://www.flowingsoftware.com)

### NbGFP in vitro infection

We obtained bone marrow derived macrophages (BMDM) by culturing bone marrow cells obtained from femur and tibia from BALB/c mice in RPMI with 30% L929-Conditioned Medium (LCM) as described by Weischenfeldt J, [27]. BMDM were co-cultured with NbGFP at M0I 1:10 for 48 hours in 6 well plates. Images were obtained and analysed using Life technologies, FLoid^®^ Cell Imaging Station.

### Statistical analysis

Intra-group and inter-group differences were compared by paired and unpaired two-tailed T-tests8.respectively. p<0.05 was considered significant. Statistical analysis was performed using Microsoft Excel.

## Acknowledgements

We are thankful for the excellent technical assistance provided by Alejandra Gallegos Velasco.Confocal microscopy was carried out under the technical supervision of Juan Carlos Segoviano Ramírez at Unidad de Bioimagen, Centro de Investigación y Desarrollo en Ciencias de la Salud (CIDICS), Universidad Autónoma de Nuevo León, Nuevo León, México

## 8 Supporting Information Captions

**Sup Fig 1.**
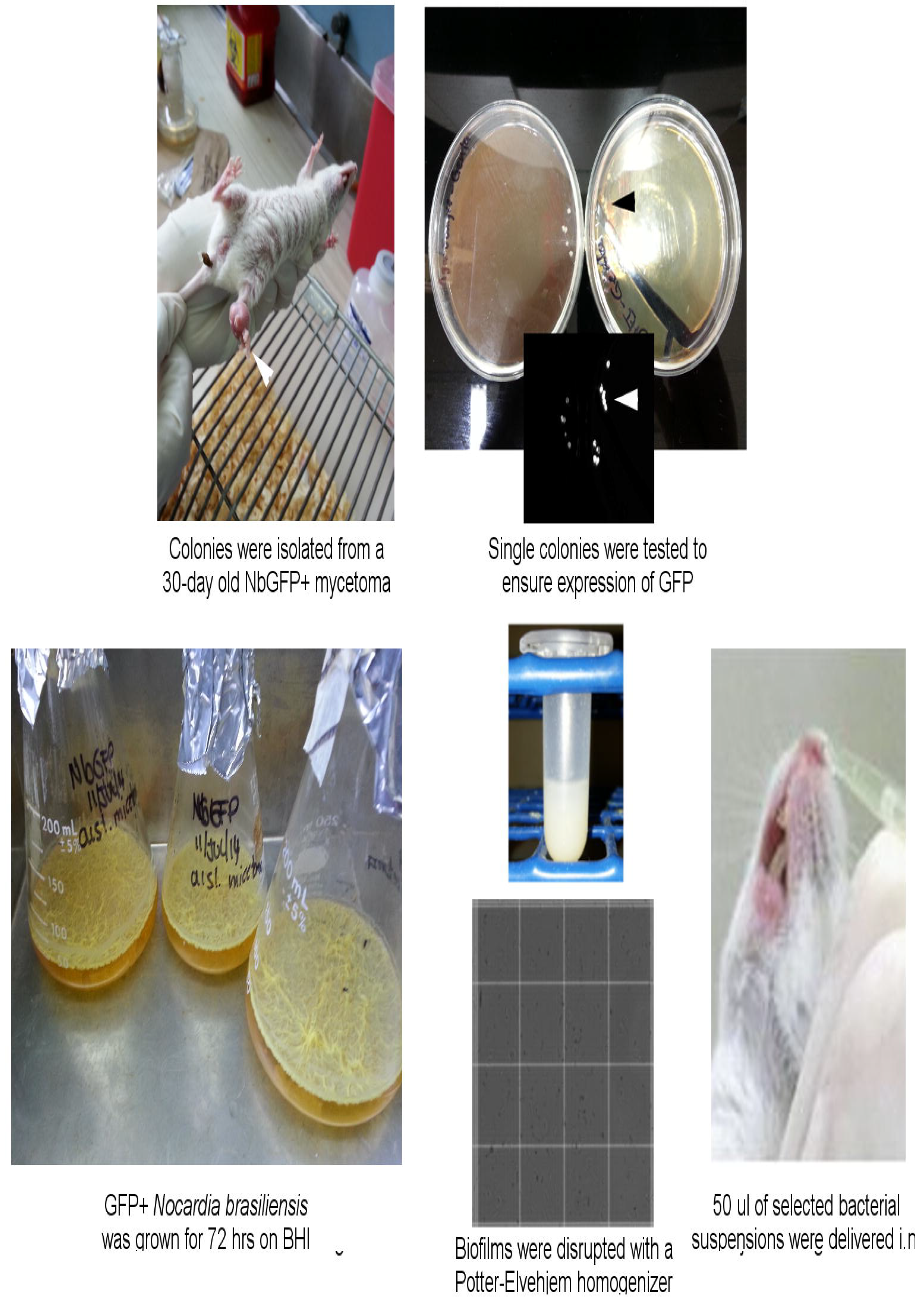
Isolation, growth and delivery of GFP+ *Nocardia brasiliensis*.

**Sup Fig 2.**
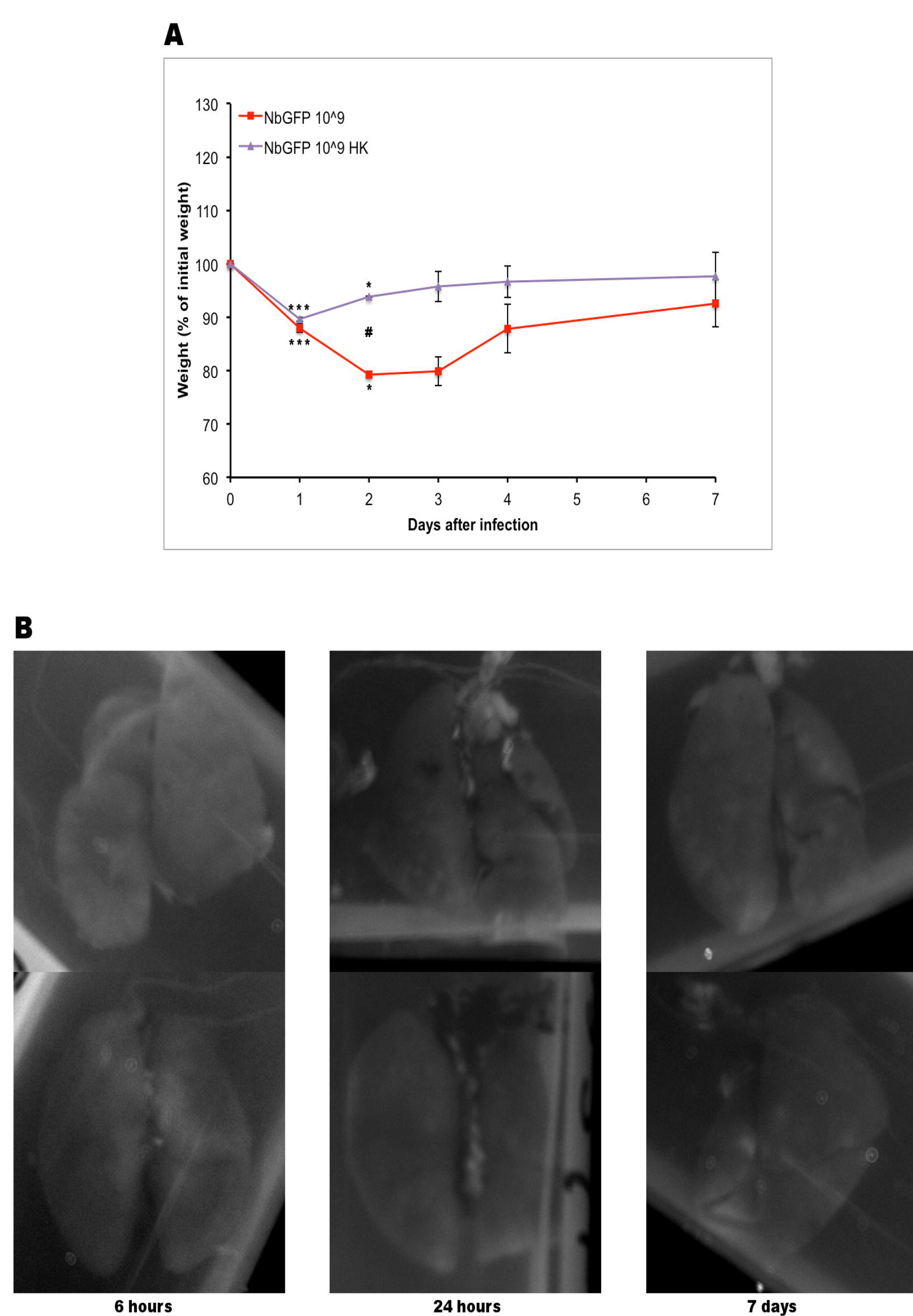
Heat killed NbGFP causes weight loss but not granulomas.

**Sup Fig 3.**
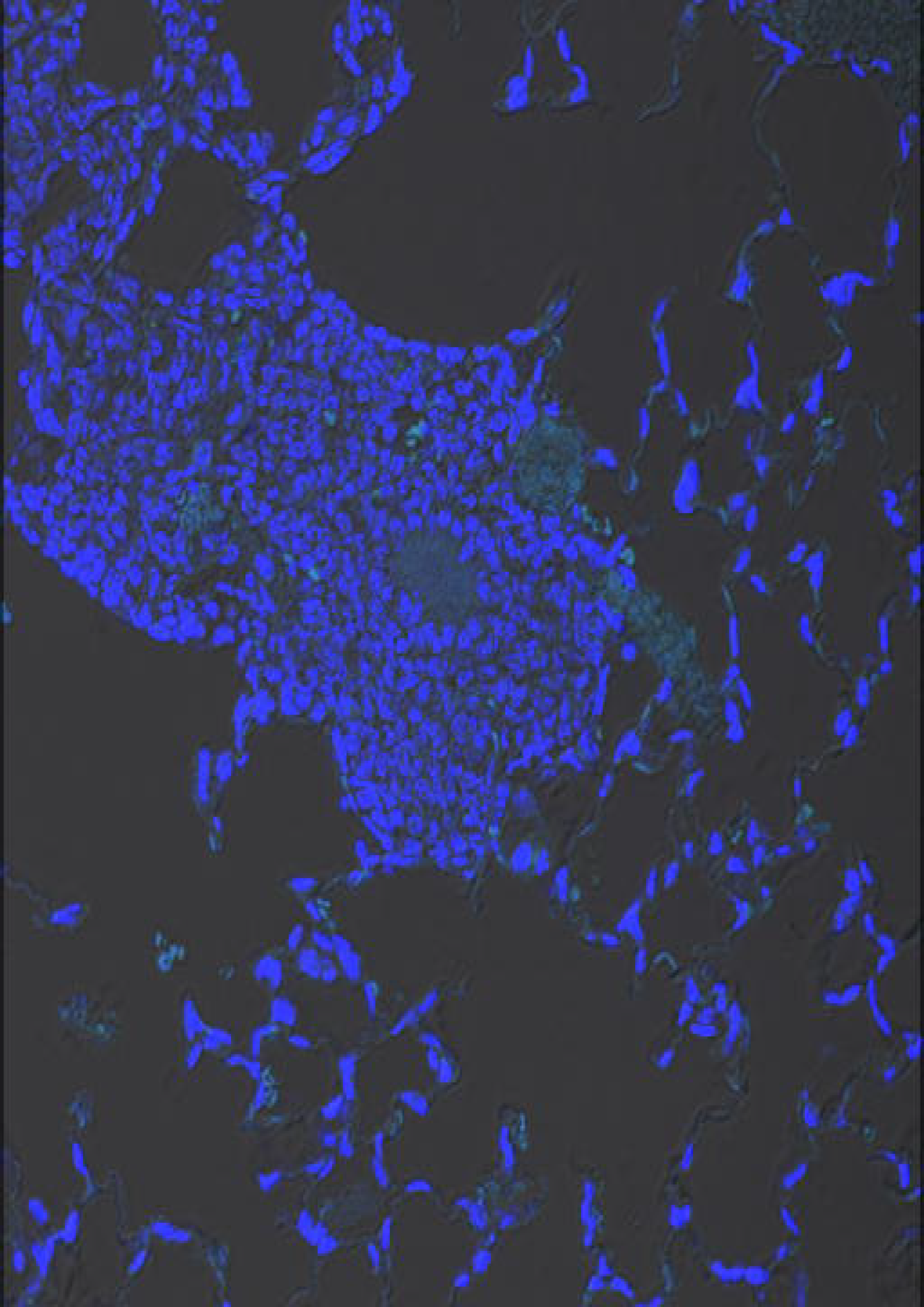
Granuloma due to wild type *N. brasiliensis*.

**Sup Fig 4.**
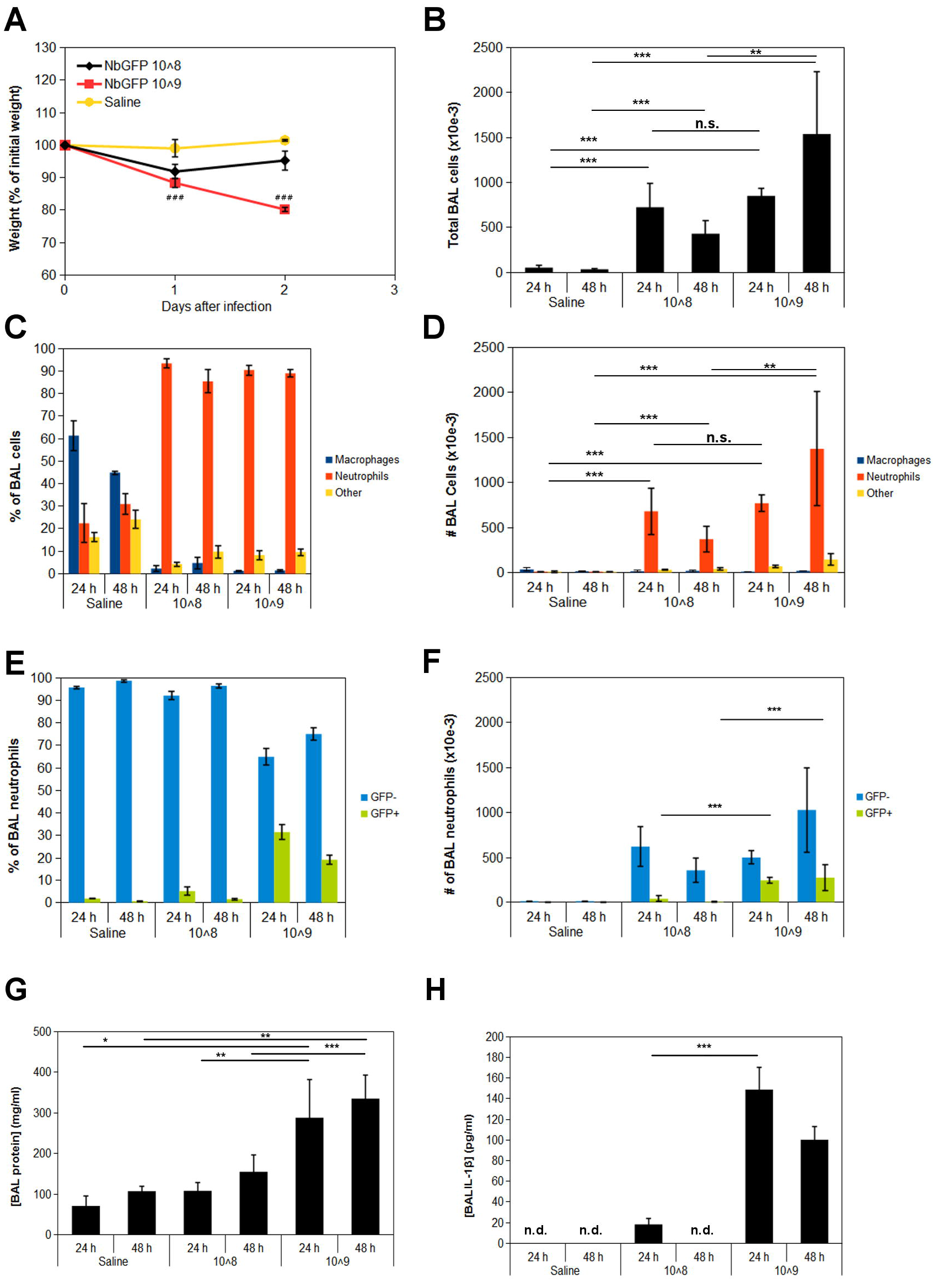
Inflammatory lung response in mice receiving either 10^8^ or 10^9^ NbGFP+ CFUs. Graphs show average +/− s.e.m. (n=5 per group) * p<0.05, ** p<0.01, *** p<0.001, ### p<0.001 10^8^ vs. 10^9^, n.d.=not detectable.

## References

1. Baracco GJ, Dickinson GM. Pulmonary Nocardiosis. Curr Infect Dis Rep. 2001;3: 286–292.

2. Menéndez R, Cordero PJ, Santos M, Gobernado M, Marco V. Pulmonary infection with Nocardia species: a report of 10 cases and review. Eur Respir J. 1997;10: 1542–1546.

3. Martínez R, Reyes S, Menéndez R. Pulmonary nocardiosis: risk factors, clinical features, diagnosis and prognosis. Curr Opin Pulm Med. 2008;14: 219–227.doi:10.1097/MCP.0b013e3282f85dd3

4. Dominguez DC, Antony SJ. Actinomyces and nocardia infections in immunocompromised and nonimmunocompromised patients. J Natl Med Assoc. 1999;91: 35–39.

5. Brown-Elliott BA, Conville P, Wallace Jr. RJ. Current Status of Nocardia Taxonomy and Recommended Identification Methods. Clin Microbiol Newsl. 2015;37: 25–32.doi:10.1016/j.clinmicnews.2015.01.007.

6. Minero MV, Marín M, Cercenado E, Rabadán PM, Bouza E, Muñoz P. Nocardiosis at the Turn of the Century: Medicine (Baltimore). 2009;88: 250–261. doi:10.1097/MD.0b013e3181afa1c8

7. Beaman BL, Goldstein E, Gershwin ME, Maslan S, Lippert W. Lung response to congenitally athymic (nude), heterozygous, and Swiss Webster mice to aerogenic and intranasal infection by Nocardia asteroides. Infect Immun. 1978;22: 867–877.

8. Ellis TN, Beaman BL. Murine polymorphonuclear neutrophils produce interferon-γ in response to pulmonary infection with Nocardia asteroides. J Leukoc Biol. 2002;72: 373–381.

9. Moore TA, Newstead MW, Strieter RM, Mehrad B, Beaman BL, Standiford TJ. Bacterial clearance and survival are dependent on CXC chemokine receptor-2 ligands in a murine model of pulmonary Nocardia asteroides infection. J Immunol Baltim Md 1950. 2000;164: 908–915.

10. Tam S, Maksaereekul S, Hyde DM, Godinez I, Beaman BL. IL-17 and γ§ T-lymphocytes play a critical role in innate immunity against Nocardia asteroides GUH-2. Microbes Infect Inst Pasteur. 2012;14: 1133–1143. doi:10.1016/j.micinf.2012.05.008

11. Mishra SK, Sandhu RS, Randhawa HS, Damodaran VN, Abraham S. Effect of Cortisone Administration on Experimental Nocardiosis. Infect Immun. 1973;7: 123–129.

12. Beaman BL. In vitro response of rabbit alveolar macrophages to infection with Nocardia asteroides. Infect Immun. 1977;15: 925–937.

13. Salinas-Carmona MC, Torres-Lopez E, Ramos AI, Licon-Trillo A, Gonzalez-Spencer D.Immune Response to Nocardia brasiliensis Antigens in an Experimental Model of Actinomycetoma in BALB/c Mice. Infect Immun. 1999;67: 2428–2432.

14. Salinas-Carmona MC, Rocha-Pizaña MR. Construction of a Nocardia brasiliensis fluorescent plasmid to study Actinomycetoma pathogenicity. Plasmid. 2011;65: 25–31. doi:10.1016/j.plasmid.2010.09.005

15. Silva CL, Faccioli LH. Tumor necrosis factor and macrophage activation are important in clearance of Nocardia brasiliensis from the livers and spleens of mice. Infect Immun. 1992;60:3566–3570.

16. Trevino-Villarreal JH, Vera-Cabrera L, Valero-Guillen PL, Salinas-Carmona MC. Nocardia brasiliensis Cell Wall Lipids Modulate Macrophage and Dendritic Responses That Favor Development of Experimental Actinomycetoma in BALB/c Mice. Infect Immun. 2012;80: 3587–3601. doi:10.1128/IAI.00446-12

17. Tam S, King DP, Beaman BL. Increase of γ§ T Lymphocyt es in Murine Lungs Occurs during Recovery from Pulmonary Infection by Nocardia asteroides. Infect Immun. 2001;69: 6165–6171. doi:10.1128/IAI.69.10.6165-6171.2001

18. Mattila JT, Ojo OO, Kepka-Lenhart D, Marino S, Kim JH, Eum SY, et al. Microenvironments in Tuberculous Granulomas Are Delineated by Distinct Populations of Macrophage Subsets and Expression of Nitric Oxide Synthase and Arginase Isoforms. J Immunol. 2013;191: 773–784. doi:10.4049/jimmunol.1300113

19. Zehra Y, Murat A, Hilal O, Mehmet AÖ, Neslihan F, Fahrettin T, and Erdoğan Ç. An Unusual Case of Pulmonary Nocardiosis in Immunocompetent Patient. Case Reports in Pulmonology. 2014. doi:10.1155/2014/963482

20. Muñoz-Hernñndez B, Noyola MC, Palma-Cortés G, Rosete DP, Galván MA, Manjarrez MEActinomycetoma in arm disseminated to lung with grains of Nocardia brasiliensis with peripheral filamentsMycopathologia. 2009 Jul;168(1):37–40. doi: 10.1007/s11046-009-9189-5. Epub 2009 Feb 24.

21. Hui CH, Au VW, Rowland K, Slavotinek JP, Gordon DL. Pulmonary nocardiosis re-visited: experience of 35 patients at diagnosis. Respir Med. 2003 Jun;97(6):709–17.doi:10.1053/rmed.2003.1505

22. Cakir E, Buyukpinarbasili N, Ziyade S, Selcuk-Duru HN, Bilgin M, Topuz U. Endobronchial nocardiosis in an 11-year-old child. Pediatr Pulmonol. 2013 Nov;48(11):1144–7. doi:10.1002/ppul.22740. Epub 2012 Dec 31.

23. Gama JA, Abby SS, Vieira-Silva S, Dionisio F, Rocha EPC. Immune Subversion and Quorum-Sensing Shape the Variation in Infectious Dose among Bacterial Pathogens. PLoS Pathog. 2012;8: e1002503. doi:10.1371/journal.ppat.1002503

24. Mihai MM, Holban AM, Giurcáneanu C, Popa LG, Oanea RM, Lazár V, et al. Microbial Biofilms: Impact 0n Pathogenesis 0f Periodontitis, CysticFibrosis, Chronic Wounds And Medical Device-Related Infections. Curr Top Med Chem. 2015;

25. Byron PR. Prediction of drug residence times in regions of the human respiratory tract following aerosol inhalation. J Pharm Sci. 1986;75: 433–438. doi:10.1002/jps.2600750502

26. Negi A, Puri A, Gupta R, Chauhan I, Nangia R, Sachdeva A. Biosafe alternative to xylene: A comparative study. J 0ral Maxillofac Pathol J0MFP. 2013;17: 363–366.doi:10.4103/0973-029X.125199

27. Weischenfeldt J, Porse B. Bone Marrow-Derived Macrophages (BMM): Isolation and Applications. CSH Protoc. 2008;2008: pdb.prot5080.

